# Mesodiencephalic junction Gabaergic inputs are processed separately from motor cortical inputs in the basilar pons

**DOI:** 10.1101/2021.10.28.466331

**Authors:** Ayoub J. Khalil, Huibert D. Mansvelder, Laurens Witter

## Abstract

The basilar pontine nuclei (bPN) receive inputs from the entire neocortex and constitute the main source of mossy fibers to the cerebellum. Despite their critical position in the cortico-cerebellar pathway, it remains unclear if and how the bPN process inputs. An important unresolved question is whether the bPN strictly receives excitatory inputs or also receives inhibitory inputs. In the present study, we identified the mesodiencephalic junction as a prominent source of GABAergic afferents to the bPN. We combined optogenetics and whole-cell patch clamp recordings and confirmed that the bPN indeed receives monosynaptic GABA inputs from this region. Furthermore, we found no evidence that these inhibitory inputs converge with motor cortex (M1) inputs at the single neuron level. We also found no evidence of any connectivity between bPN neurons, suggesting the absence of a local circuit. Finally, rabies tracings revealed that GABAergic MDJ neurons themselves receive prominent inputs from neocortical output neurons. Our data indicates that inhibition from the MDJ, and excitation from the neocortex remain separate streams of information through the bPN. It is therefore unlikely that inhibition in the bPN has a gating function, but rather shapes an appropriate output of the bPN during behavior.

## Introduction

Motor control relies on brain-wide networks. Motor cortex directs voluntary movements (Guo et al. 2015) and the cerebellum coordinates movements (Manto et al. 2012). Reciprocal connections between these structures are necessary for proper motor control. Indeed, the cerebellum projects to motor cortex via the thalamus (Sawyer et al. 1994; Aumann 2002; Gornati et al. 2018), while motor cortex projects to the cerebellum via the pontine nuclei (Schwarz and Mock 2001; Kratochwil et al. 2017). This closed-loop connectivity enables forward and inverse models for motor control (Wolpert et al. 1998; Shadmehr and Krakauer 2008). Interestingly, other parts of neocortex and cerebellum are also connected (Kelly and Strick 2003; Henschke and Pakan 2020; Pisano et al. 2021), enabling similar computational mechanisms for motor control and cognitive processes alike.

This places the pontine nuclei at the nexus of information transfer between neocorex and cerebellum. Indeed, the mossy fiber afferents from the basilar pontine nuclei (bPN) to the cerebellum is one of the largest fiber tracts in the brain Additionally, the bPN also receives inputs from numerous non-neocortical regions of the brain (Burne et al. 1981; Wiesendanger and Wiesendanger 1982; Kosinski et al. 1986; Mihailoff et al. 1988, 1989). These non-cortical and corticopontine afferents show a topographical organization with minimal regional overlap within the bPN (Leergaard and Bjaalie 2007; Proville et al. 2014; Kratochwil et al. 2017). Similarly, mossy fibers originating from the bPN project to specific zones in the cerebellum (Päällysaho et al. 1991; Mihailoff 1993; Odeh et al. 2005; Huang et al. 2013; Kratochwil et al. 2017). Therefore, the bPN is often not considered to have an active role in integration of information, but is often considered to be a relay for information destined for the cerebellum.

Still, synaptic plasticity of inputs to the bPN has been described, suggesting a potential way of processing of inputs to bPN neurons (Möck et al. 1997), potentially shaping spiking activity in the bPN (Schwarz et al. 1997; Möck et al. 2006; Guo et al. 2021). Additionally, various extrinsic sources of inhibition to bPN neurons have been suggested (Border et al. 1986; Mihailoff and Border 1990; Möck et al. 1999), but these sources of GABAergic inputs to the bPN have never been physiologically confirmed or characterized, precluding conclusions about their function and integration in the cerebro-cerebellar circuit.

Here we identify the mesodiencephalic junction (MDJ) as the main source of GABAergic signaling to the bPN. This inhibition does not seem to interact with afferents from motor cortex, even though their projections overlap in the bPN. In contrast to the strongly depressing motor cortex inputs, GABAergic inputs from MDJ show remarkably little short-term depression. Finally, using rabies-tracing we show that pontine-projecting MDJ neurons receive prominent neocortical inputs, similar to bPN neurons themselves. These results suggest that the bPN contains separate streams for processing information from neocortex directly, and sign-inverted neocortical inputs.

## Materials & Methods

### Animals

Male and female wt C57BL/6J mice were used for acute slice experiments. Animals were housed socially (max. four per cage) and had ad libitum access to chow and water. All experimental procedures were approved by the Central Authority for Scientific Procedures on Animals and local animal welfare body of the VU University and VU University Medical Center (Amsterdam, Netherlands) and carried out in accordance with European and Dutch law.

### Intracranial virus and tracer injections

Microinjection needles were pulled from 3.5” borosilicate glass capillaries (Drummond SCI, USA) on a Sutter P-87 puller (Sutter, CA) and backfilled with mineral oil before virus solution was loaded. AAV9 viruses were purchased from Addgene (USA) syn.Chronos-GFP.WPRE.bGH and syn.ChrimsonR-tdTomato.WPRE.bGH were injected at 4·10^12^ vg/ml titer and 1.5·10^12^ vg/ml respectively. Retrograde AAV2 virus was purchased from University of Zurich vector core. AAV2r-hSyn1-chI-iCre-WPRE-SV40 was injected at a titer of 7.9·10^12^ vg/ml. Rabies virus (Rabies-SAD-dG-tdTomato) and AAV helper virus (rAAVdj-hsyn1-dlox-TVA-2A-EGFP-2a-oG(rev)-dlox-WPRE-bGhp(A)) were a generous gift from Klaus Conzelmann. All mice used for optogenetic experiments received intracranial virus injections at postnatal 21. For all surgeries, mice received Carprofen (5 mg/kg s.c.) and Buprenorphine (50 μg/kg s.c.) pre-operatively. A second Carprofen injection (5 mg/kg s.c.) was administered twenty-four hours post-surgery. Animals were kept under general anesthesia during surgery with Isoflurane (0.5% - 1%). Ear bars were placed to secure the skull, a small amount of Lidocaine cream was applied before placement. Local analgesia was applied by injecting a small volume of Lidocaine (2%) underneath the scalp before incising the skin. The scalp was cut and folded open to expose the skull, holes were drilled to access the injection sites, and virus was delivered via injection. (relative to bregma(Paxinos and Watson 1998) (in mm), M1: AP 1.30; ML 1.08L; DV 1.20, MDJ: AP -3.50; ML 0.50L; DV 3.00 Cerebellum: AP -6.2; ML 1.5R; DV 2.0 bPN: AP-4.0 ML 0.5L DV 5.5). For optogenetic experiments, total volume of 500 nl was injected per site in steps of 50 nl/min using a Nanoject II (Drummond SCI, USA) set to the ‘slow’ rate (23 nl/sec). The microinjection needle was left in place for 5 minutes before and after injection. Mice were sacrificed for acute slice experiments at least two weeks after viral injection to allow for adequate expression. For tracing experiments, total volumes between 10 and 100nl were injected per site and needles were left in place for 15 minutes before retraction. Retrobead transport was assessed after 14 days. For rabies tracing animals were injected with AAV to express cre, oG and TVA in one surgery. After 1 week rabies virus was injected, after which we waited another week before animals were perfused with 4% formaldehyde solution in 0.1M phosphate-buffered saline (PBS) for analysis.

### Acute slice preparation

Acute slices were prepared for optogenetic experiments (sagittal orientation) and paired recordings (sagittal or coronal). Before decapitation, mice first received a lethal pentobarbital injection (120 mg/kg i.p.) and were perfused with ice cold N-Methyl- D -glucamine (NMDG) solution containing (in mM): NMDG 93, KCl 2.5, NaH_2_PO_4_ 1.2, NaHCO_3_ 30, HEPES 20, Glucose 25, sodium pyruvate 3, sodium ascorbate 5, MgSO_4_ 10, CaCl_2_ 0.5, adjusted to 315 mOsm ± 5 and pH 7.3. After decapitation, the brain was removed from the skull and sliced in the same oxygenated ice-cold NMDG solution. Brains were sliced using a ceramic blade (Campden Instruments ltd., England) and slices (250 μm) were collected in an oxygen-perfused brain slice chamber filled with a holding solution containing (in mM): NaCl 92, KCl 2.5, NaH_2_PO_4_ 1.2, NaHCO_3_ 30, HEPES 20, Glucose 25, sodium pyruvate 3, sodium ascorbate 5, MgSO_4_ 10, CaCl_2_ 0.5, adjusted to 305 ± 5 mOsm. Slices were kept oxygenated at room temperature until the moment of recording.

### Acute slice whole-cell recordings

During all acute slice experiments, whole-cell recordings were acquired at a temperature of 33 ± 1 ^□^C. Brain slices were placed in a bath continuously perfused with oxygenated ACSF containing (in mM): NaCl 125, KCl 2.5, NaH_2_PO_4_ 1.25, NaHCO_3_ 26, Glucose 25, MgCl_2_ 1, CaCl_2_ 1.3, adjusted to 305 ± 5 mOsm. Borosilicate glass capillaries were pulled to produce patch-pipettes with a resistance of 3 – 6 MΩ. For optogenetic experiments, patch-pipettes were filled with a cesium methanesulfonate-based pipette solution containing (in mM): CsMethanesulfonate 115, TEA 25, HEPES 10, EGTA 0.2, QX-314 Cl 5, NaCl 4, MgATP 2, Na_3_GTP 0.4, Na_2_Phosphocreatine 10. For paired patch-clamp experiments, patch-pipettes were filled with a potassium gluconate-based solution containing (in mM): KGluconate 135, KOH 31, NaCl 10, HEPES 10, EGTA 10, Na_2_ATP 4, Na_3_GTP 0.4. Both internal solutions were adjusted to pH 7.2 and 310 mOsm. Biocytin (0.05%) was added to internal solution on the day of the experiment. Cells were loaded with biocytin during whole-cell patch clamp recordings and resealed at the end of the experiment. Slices were then transferred to paraformaldehyde (PFA, 4%) and fixed for at least 48 hours.

### Optogenetic stimulation

Optogenetic responses were evoked using a 4-channel LED system (DC4100 & LED4D114; Thorlabs inc., USA). Cells were voltage clamped at -70 and 0mV and screened for responses using full field 100 ms optical stimulation at all four wavelengths (405, 470, 505 & 590nm). When responses (EPSCs or IPSCs) were observed, the light source was restricted to a small beam (±100μm diameter) with high intensity (>100mW/mm^2^)(Jackman et al. 2014) to allow reliable axonal stimulation of afferents. Optical stimulation was delivered in trains of twenty pulses with a 10 second intertrain interval, and was alternated per sweep in a pseudorandom order (20 – 100 – 50 – 10Hz). All input characterizations are based on afferents expressing Chronos. A short negative voltage (50 ms, -10 mV) was injected at the start of each sweep to monitor access resistance throughout the experiment. Voltage clamp recordings were acquired at a 50.0 kHz sample rate with a 10 kHz low pass filter. Cells from optogenetic experiments were analyzed on the following conditions: (1) optical stimulation at 470 or 590 nm evoked a response at -70 mV or 0 mV holding potential; (2) at least nine sweeps per frequency were collected. Responses following stimulation were defined as optogenetically evoked inputs if they exceeded the threshold set at 2σ of the baseline. Responses that did not reach the computed threshold were not considered in the analysis. Response amplitudes were computed on averaged sweeps. The peak amplitude was detected within an eight-millisecond time window after each light pulse. Then, the response was determined by calculating the average maximum amplitude over a one-millisecond time window of the peak amplitude. Baseline was defined as the average amplitude over a two-millisecond time window before optic stimulation.

### Paired recording

Sagittal or coronal slices were prepared for paired recordings. Up to three neurons were recorded at the same time, and potential connections between neurons were probed by evoking spike trains successively in each neuron. Ten action potentials were evoked presynaptically using current injections of 2nA at 50Hz, followed by a single current injection after 500 ms. Cells were kept at or around resting membrane potential throughout recording to detect EPCSs. Current clamp recordings were acquired at a 50.0 kHz sample rate with a 10 kHz low pass filter. We did not compensate for the liquid junction potential.

Cells from paired whole-cell patch clamp experiments were analyzed when: (1) stimulation evoked action potentials (APs); (2) cells did not have a negative leak current exceeding 500 pA; (3) recordings had a stable resting membrane potential; (4) at least fifteen sweeps were collected. To detect connections, we looked for EPSCs in the average postsynaptic response in the first 18ms after the AP to accommodate for mono- and disynaptic connections. Then, the postsynaptic response was determined by calculating the average amplitude over a one-millisecond time window of the peak amplitude.

### Pharmacology

Gabazine (10 μM) was bath-applied to inhibit post-synaptic GABA_A_ responses. AMPA receptors were inhibited with 6,7-dinitroquinoxaline-2,3-dione (DNQX, 10 μM). Glycine receptors were inhibited by application of Strychnine (1 μM). TTX (1 μM) was applied to inhibit voltage-gated sodium channels. Voltage-gated potassium channels were inhibited with 4-Aminopyridine (4-AP, 100 μM). All antagonists were bath-applied and perfused at least five minutes before the start of a recording.

### Histology

For neurons recorded in vitro, slices were first washed in phosphate-buffered saline (PBS, 0.1 M, 3×15’) containing (in mM): NaCl 137, KCl 2.7, NaH_2_PO_4_ 12, KH_2_PO_4_ 1.8. Then, slices were permeabilized in Triton-X (PBS-T, 0.5%; 2h). Following permeabilization, slices were again washed in PBS (3×15’) and stained with Streptavidin-Alexa 647 (1:500 in 0.5% PBS-T). Finally, slices were washed in PBS (4×15’) and embedded in Mowiol (2%) on glass microscope slides for confocal imaging. Tracer-injected brains were washed in 0.1M PBS and embedded in 11% gelatin for whole-brain sectioning. 50-100μm sections were made on a Leica VS1000 vibratome and collected directly to glass slides (retrobead tracing, coronal slices), or in 3 jars per side (sagittal slices, 6 jars total per brain). Staining for GAD was performed on one jar per side of the brain, yielding 24-30 sections for analysis. Sections were washed 3×5’ in PBS with 0.025% Triton-X, blocked for 30’ (PBS+0.025%TX and 5% normal Donkey Serum), and subsequently incubated with GAD67 antibody O/N (1:500 MAB5406, Merck-Millipore) in PBS. The next day sections were washed 3×5’ in PBS, incubated for 2 hours with Alexa647 secondary (Goat anti mouse, A21235, ThermoFisher), again washed 3×5’ in PBS and mounted on slides with mowiol (2%). Cells recovered from in vitro recordings, and sections from rabies tracing experiments were imaged on a confocal microscope (Nikon), retrobead tracing was visualized on a epifluorescence microscope (Zeiss). Retrobead tracing was analyzed by first marking all labeled neurons via cellcounter in matlab (https://github.molgen.mpg.de/MPIBR/CellCounter) and then aligning sections while the wholebrain tool in R (Fürth et al. 2018). Sections between B+3.0 and B-5.6 were aligned to the Allen Brain Atlas.

All display items show mean ± SEM unless noted otherwise.

## Results

To anatomically reveal cortico-cerebellar pathways that run via pontine nuclei, we first injected retrobeads into the cerebellum in p21 mice (**Figure 1**). Injections were done in the white matter of paravermal lobule 5 (**Figure 1A,** N=2) and retrograde labeling was photographed after 14 days. As expected, retrograde labeling from the cerebellum could be observed in the inferior olivary nucleus, external cuneate nucleus, lateral reticular nucleus and basilar pontine nucleus (**Figure 1B**), but labeling was completely absent from cerebral cortex (**Figure 1B**). To investigate inputs to the basilar pontine nuclei, we injected retrobeads into the basilar pontine nuclei of p21 mice (**Figure 1C,** N=2). Injections were confined to the basilar pons, with minimal invasion of overlying structures (**Supplementary Figure 1**). We observed predominantly inputs from the ipsilateral side of the brain (**Figure 1G**), with almost all labeling (90% of retrogradely labeled neurons) in deep layers (**Figure 1F**) throughout the neocortex (**Figure 1E**). Of other areas providing afferents to the bPN midbrain was most prominent (5%, with the mesodiencephalic junction, MDJ, being particularly prominent with 3% of brain-wide retrobead signal, **Figure 1H**), followed by thalamus and hypothalamus (3%), **Figure 1E**).

**Figure 1:**
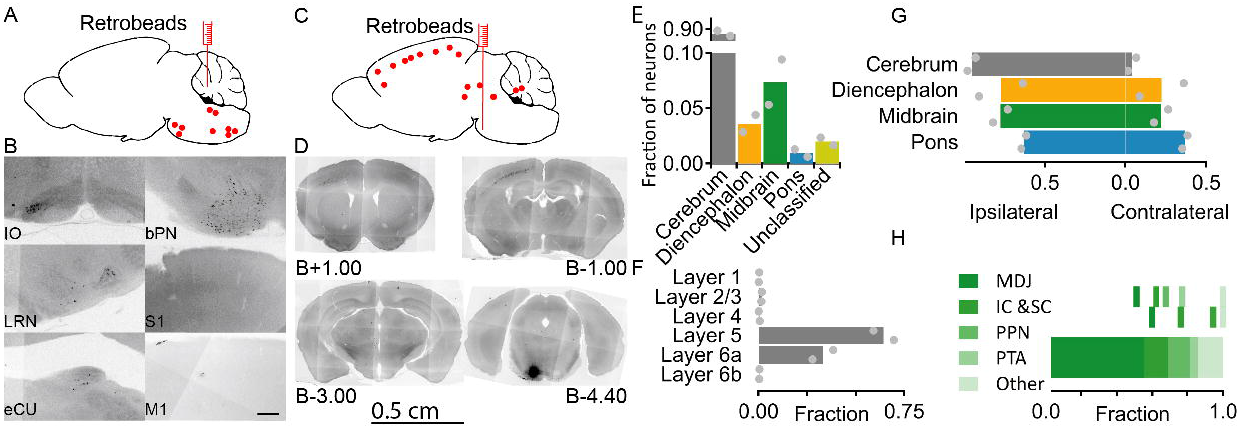
The basilar pontine nuclei are intermediate between the cerebral cortex and the cerebellum. (A) Schematic representation of a retrobead injection in cerebellum. (B) Examples of retrobead-labeled neurons in the inferior olivary nucleus (IO), lateral reticular nucleus (LRN), external cuneate nucleus (eCU), basilar pontine nuclei (bPN), and the absence of labeled neurons in primary sensory cortex (S1), and primary motor cortex (S1). Scale bar represents 50 μm. (C) Schematic representation of a retrobead injection in the bPN. (D) Example retrograde labeling after injections in bPN. Scale bar represents 500 μm. (E) Quantification of retrograde labeling in cerebrum, diencephalon, midbrain, pons and other areas. Shown is the average and individual data points. (F) Retrograde labeling in neocortex is predominantly found in deeper layers (layer 5 and 6a), consistent with the location of extratelencephalic projection neurons. (G) Retrograde labeling can be predominantly found ipsilateral. (H) Of the prominent inputs from midbrain, MDJ provides the most prominent input to bPN.

To characterize short-term plasticity of cortico-pontine synapses, we injected AAV to drive expression of Chronos-GFP in the M1 region to enable visualization and stimulation of M1-specific afferents in the bPN. Whole cell recordings from bPN neurons were made in sagittal slices, in which axons from M1 were stimulated with short pulses of blue light focused in a small spot (100 μm diameter) >500μm away from the soma of the recorded neuron (**Figure 2A,B**)(Jackman et al. 2014). All inputs evoked from motor cortex had a short delay to onset (2.4 ± 0.06 ms), a fast rising phase and decay (10-90%: 1.0 ± 0.22 ms and 90-10%: 20 ± 12 ms, respectively), and were reduced to 4.4 ± 0.8% of the original response by the AMPA and kainate receptor antagonist DNQX (10 μM; ACSF: 20 ± 5. pA; DNQX: 1.0 ± 0.3 pA; p<0.001; n = 4 neurons; **Figure 2C**), confirming that neocortical inputs to bPN are glutamatergic. During train stimulation we observed prominent short-term synaptic depression of M1 inputs across all frequencies, with more pronounced depression at higher frequencies and later in the stimulus train (**Figure 2E,** n=4 cells in n=4 animals). To check for possible opsin-specific influences on midbrain inputs, we repeated these experiments with ChrimsonR instead of Chronos expressed in M1. In these experiment we noticed that at higher stimulation frequencies Chrimson-evoked responses were more depressed after a train of stimuli (**Supplementary Figure 2**). Since this is likely due to incomplete recovery of the channelrhodopsin variant (Klapoetke et al. 2014), we continued our characterization of short-term plasticity of cortico-pontine synapses using Chronos. In animals in which Chronos was expressed in motor cortex, synaptic responses recorded in the bPN were attenuated at 50 and 100Hz after a 20-pulse train stimulus to 0.7 ± 0% and 0 ± 0% of initial amplitude respectively (steady-state, average of last five responses). After train stimuli at 10 and 20Hz, responses were attenuated to 41 ± 3% and 22 ± 2% respectively. To confirm that neocortex makes monosynaptic connections to the bPN, we applied the voltage-gated sodium channel blocker tetrodotoxin (TTX) to block AP-generated neurotransmitter release, followed by the combined application of TTX and 4-aminopyridine (4-AP). As expected, optogenetically-evoked responses were virtually absent in the presence of TTX (reduced to 2.2 ± .2% of the original response; ACSF: 50 ± 18 pA; TTX: 1.0 ± 0.7 pA; n=3; p<0.001. Subsequent addition of 4-AP, which blocks voltage-gated potassium channels and prolongs optogenetically-evoked depolarization, recovered the synaptic responses (130 ± 56% of amplitude in ACSF; TTX + 4-AP: 50± 28 pA, n=3; p=0.66 vs ACSF; **Figure 2D**). These results show that motor cortex provides prominent, but strongly depressing monosynaptic glutamatergic inputs to bPN neurons.

**Figure 2:**
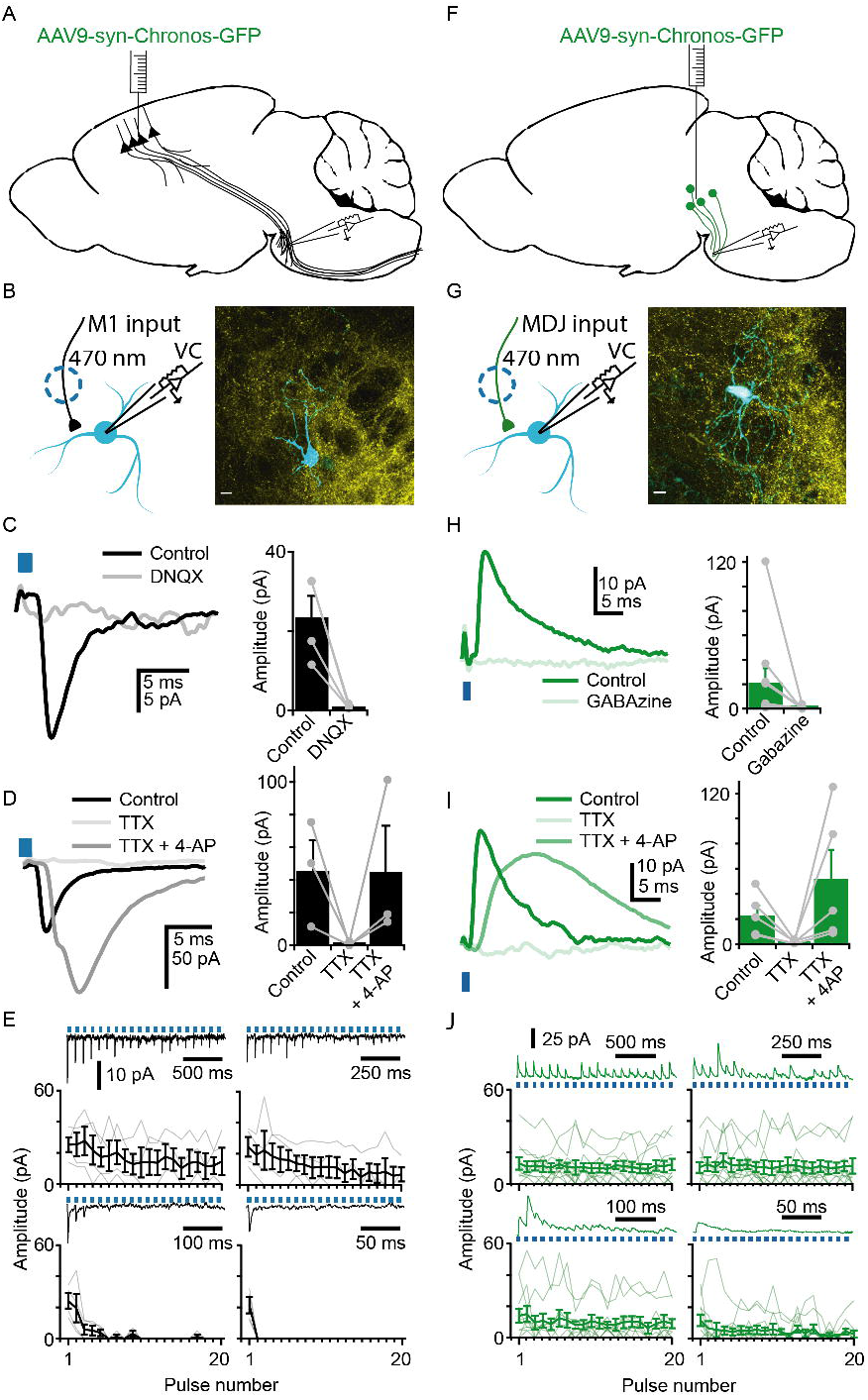
Optogenetic stimulation of neocortical and mesodiencephalic afferents to the bPN. (A) Schematic overview of intracranial virus injection in M1. (B) Schematic overview of the experimental patch-clamp approach (left), and post-hoc recovered and stained neuron (cyan) showing GFP+ fibers from M1 (Yellow). (C) neocortical inputs can be effectively blocked with an antagonist against the AMPA and kainate receptor. Example trace (left) and quantification (right). (D) Neocortex makes monosynaptic contacts to bPN. Example trace (left) and quantification (right). (E) Neocortical inputs are strongly depressing, especially at high frequencies. (F) Schematic overview of intracranial virus injection in MDJ. (G) Schematic overview of experimental approach (left) and a recovered neuron (cyan) with GFP+ fibers from MDJ (Yellow) (H) MDJ inputs can be effectively blocked with an antagonist against the GABA_A_ receptor. Example trace (left) and quantification (right). (I) MDJ makes monosynaptic contacts to bPN. Example trace (left) and quantification (right). (J) MDJ inputs show very limited short-term depression, even at higher frequenies.

Our retrograde tracing experiments (**Figure 1**) have shown that in addition to several neocortical regions, several subcortical brain regions provide inputs to the bPN. These subcortical brain regions might provide non-glutamatergic inputs, as has been suggested before (Border et al. 1986; Mihailoff et al. 1988, 1989; Mihailoff 1995). Furthermore, several studies have consistently reported the presence of GABAergic boutons in the bPN, suggesting a possible role of inhibitory signaling (Mihailoff and Border 1990). However, the exact source and potential role of these inputs has not been elucidated, nor electrophysiologically isolated and characterized. To investigate this in the bPN, we stained sections of mouse brain using an antibody against the enzyme Glutamate decarboxylate 67 (GAD67), which is present in somata and synapses of GABAergic neurons. In these animals we never observed GAD67+ somata, but we did observe prominent and numerous GAD67+ boutons (**Supplementary Figure 3A,B**; N = 6 mice). Similarly, in GAD-GFP mice (Chattopadhyaya et al. 2004), we never observed GFP+ somata, but we observed putative axons stained for GFP in the bPN (**Supplementary Figure 3C**; N = 4 mice). This indicates that there is a prominent extranuclear source of GABA to the bPN. Closer investigation of the afferent areas in midbrain revealed that the majority of inputs from midbrain arose from the mesodiencephalic junction (MDJ) (3% of all projections to the bPNm, **Figure 1H**), an area intimately involved with the cerebellar circuit (Ruigrok 2004). Even though glutamatergic neurons of the MDJ project to the inferior olive (de Zeeuw et al. 1989; Ruigrok and Voogd 1995), these neurons are positioned intermixed with neurons that contain other neurotransmitters (De Zeeuw and Ruigrok 1994).

To confirm that the MDJ is the source of GABAergic inhibition to the bPN, we injected AAV to express Chronos-GFP in this region. In acute slices we performed whole cell recordings in the bPN in the area that receives afferents from motor cortex. In these experiments we observed prominent outward currents in neurons held at 0 mV, with a short rise, and long decay (2.1 ± 0.36 ms and 140 ± 46 ms, respectively) when stimulating with light (**Figure 2F**). These inputs were reduced to 9 ± 19% in the presence of the GABA receptor antagonist Gabazine (ACSF: 20 pA ± 11 pA; Gabazine: 0.5 ± 0.41 pA; n = 10 neurons; p < 0.001 ; **Figure 2H**). But, we did not observe a change in holding current (ACSF: 140 ± 28 pA vs Gabazine: 160 ± 33 pA, n=10 neurons, p=0.27), indicating that the inhibition from MDJ and in bPN neurons is predominantly phasic.

In contrast to glutamatergic synaptic inputs from neocortex, GABAergic synaptic input from MDJ neurons showed remarkably little short-term synaptic plasticity with intervals >20 ms, even after a stimulation-train of 20 pulses we observed 108 ± 5% and 105 ± 7% of the initial amplitude for 10Hz and 20Hz stimulation trains respectively (**Figure 2J**). Only at frequencies ≧50Hz and towards the end of a train of pulses was the amplitude of responses depressed (to 55 ± 7% and 14 ± 7% for 50 and 100Hz stimulus trains, respectively). Similar to responses evoked from M1 afferents, MDJ afferents showed enhanced short-term synaptic depression when we evoked responses from efferents that expressed the channelrhodopsin variant Chrimson (**Supplementary Figure 2**). To confirm that inputs from the MDJ are monosynaptic, we applied TTX followed by the combined application of TTX and 4-AP. Inputs from the MDJ are completely blocked upon TTX application (5 ± 3.9% of the response in ACSF; ACSF: 23 ± 8 pA vs TTX: 1.5 ± 0.61 pA; p < 0.001, n = 5 neurons) and subsequently rescued in presence of 4-AP (to 400 ± 760% of ACSF response; TTX +4-AP: 50 ± 23pA; p = 0.37, n = 5 neurons **Figure 2I**).

Our results thus far indicate the neurons in the bPN receive depressing excitatory input from neocortex and inhibitory input from the MDJ that undergoes very little short-term plasticity. These inputs could interact in several ways in the cerebro-cerebellar circuitry. First we investigated whether single neurons in the bPN receive inputs from both motor cortex and from MDJ. We therefore analyzed full field optical stimulation data of all neurons that received inputs from either M1 or MDJ (See methods). Neurons were clamped at -70 mV and subsequently at 0 mV to enable detection of both EPSCs and IPSCs, respectively, in the same neurons (**Figure 3A-C**). Out of 53 recorded bPN neurons that responded to optogenetic stimulation, 62.26% (33 out of 53) of neurons only received inputs from MDJ and 37.73% (20 out of 53) only received inputs from M1 **Figure 3D**). No neurons that responded to optic stimulation received inputs from both M1 and MDJ, suggesting that these afferents make synapses onto different neuron classes within the bPN. To test this, we compared several passive electrical properties between the two groups. However, we found no statistically significant differences in membrane resistance (M1: 320 ± 49 MΩ; midbrain: 220 ± 25 MΩ; p = 0.08; n = 52 neurons), membrane capacitance (M1: 100 ± 15 pF; MDJ: 108 ± 8.2; p = 0.86; n = 52 neurons) and membrane decay time constant (M1: 1.18 ± 0.08 ms; MDJ: 1.2 ±0.10 ms; p = 0.96; n = 52 neurons) between these two groups, indicating that these neurons receive different inputs but do not represent separate classes of neurons (**Supplementary Figure 4**). Thus, our results show that convergence of inputs from M1 and MDJ in the bPN is at best remarkably rare. This overall segregation of excitatory and inhibitory streams suggests that it is unlikely that MDJ inputs modulate incoming motor inputs to bPN neurons.

**Figure 3:**
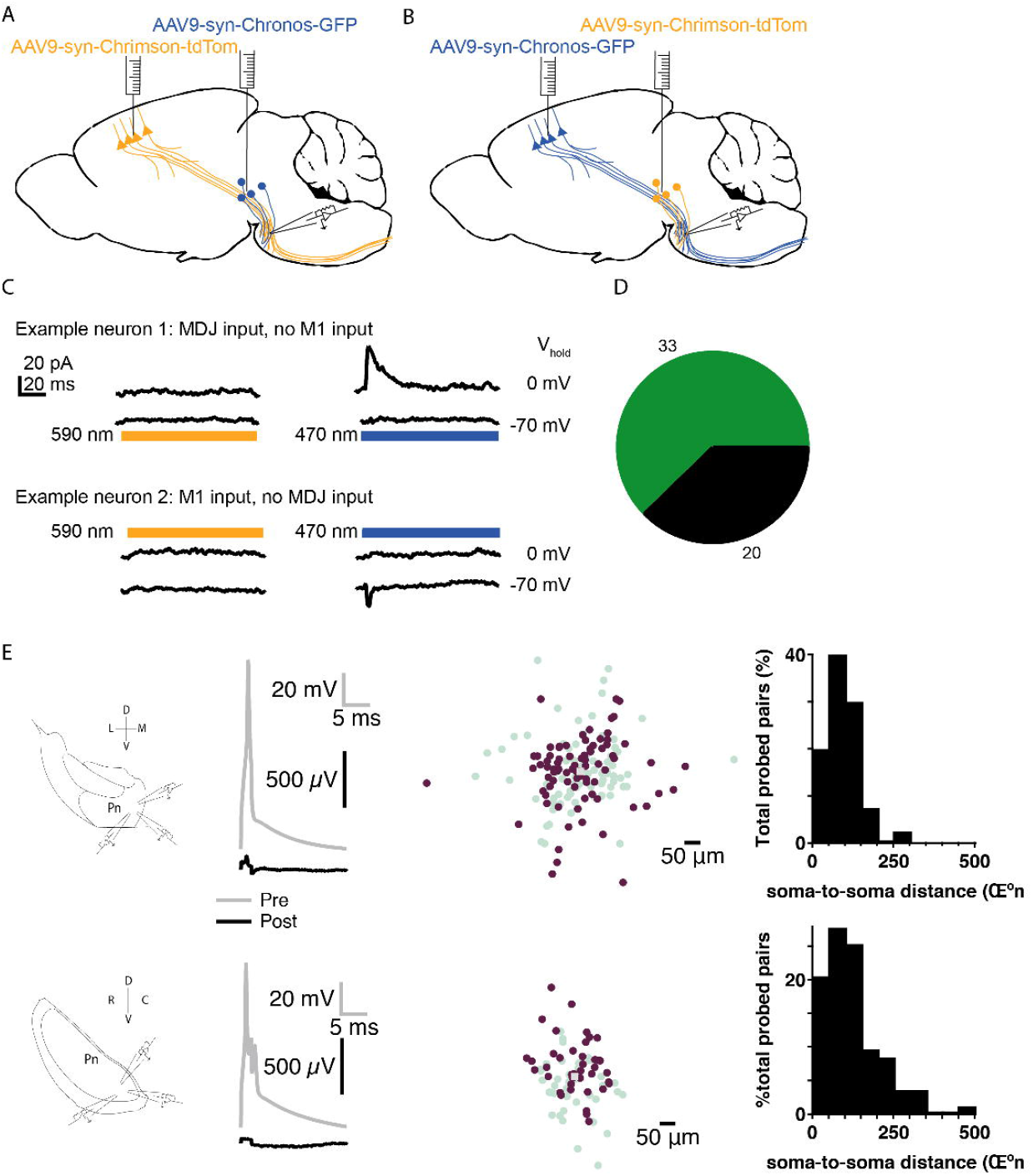
The bPN receives monosynaptic GABAergic inputs from midbrain. (A) Schematic overview of intracranial virus injection configuration in midbrain (Chronos) and M1 (Chrimson) (left), or inverse (right) (B) Single neurons either receive input from neocortex, or from mesodiencephalic junction, but not from both. Shown are two example neurons respoinsive to blue light, but not to yellow light. (D) Quantification of inputs from neocortex and from mesodiencephalic junction. (E) Experimental setup whole-cell patch clamp recordings in coronal (top) and sagittal (bottom) sections of bPN with example paired recordings of bPN neurons in coronal slice (top) and sagittal slice (bottom). Presynaptic neurons are indicated in black, postsynaptic neurons in grey. Right: Distances of probed connections measured between pre-synaptic neuron (black square, middle) and post-synaptic neuron (purple). Teal markers indicate reciprocal distances and are point-mirrored to purple markers. Dorsoventral and lateromedial orientation as shown in (A). Far right: Soma-to-soma distance distribution of all probed pairs for coronal (top) and sagittal (bottom) experiments. D: Dorsal; V: Ventral; L: Lateral; M: Medial.

Nonetheless, there is a distinct possibility that inputs from M1 and MDJ can interact in the MDJ via a local network between bPN neurons. One study has suggested that short-range interactions between neurons in the bPN are absent, but this dataset only comprised of 20 tested pairs and only probed connections within a short range from each other (Möck, Schwarz, 2006). To investigate this issue over a longer distance, we recorded from a total 125 pairs (i.e. 250 unidirectional connections) spaced up to 500μm apart, in slices cut in both the coronal (n = 168) and sagittal (n = 82) orientation to avoid confounding effects of slice orientation (Shinoda et al. 1992) (**Figure 3E**). In these sampled pairs we never detected evidence of mono- or disynaptic contacts between neurons. Therefore, it is unlikely that M1 and MDJ inputs interact at the level of either single neurons or within a local circuit, but rather that M1 and MDJ inputs are processed separately

We have thus far shown that bPN neurons receive GABAergic inputs from the MDJ and that at the level of the bPN these inputs do not interact with inputs from M1. Furthermore, this inhibition is predominantly mediated phasically rather than via a tonic current. Therefore, it is unlikely that input from the MDJ induce a dampening or level-setting effect on neurons of the bPN (Silver 2010). Another possible role for inhibition is the gating of information (Crowley et al. 2009; Geborek et al. 2013). However, in that case one would expect that inputs interact at the level of the bPN, which is not in line with our data. A final possibility is that input from the MDJ represents a separate stream of information through the pons to the cerebellum. This possibility could explain recent reports that distinct pontine neuron populations either increase or decrease their firing rates during a voluntary reaching and grabbing task (Guo et al. 2021). If bPN-projecting MDJ neurons are indeed recruited during movement, we expect that they receive prominent inputs from the neocortex. We investigated this possibility by using monosynaptic rabies tracing (Wickersham et al. 2006).

To map inputs to neurons in the MDJ that provide inputs to the bPN, we first checked whether we could trace connections from neocortex, through bPN to cerebellum. Indeed, by injecting a retrograde virus into cerebellum to express cre in bPN neurons, followed by viruses to express TVA and rabies glycoprotein in bPN, we could visualize rabies-infected neurons in neocortex after injections with rabies virus in bPN (**Supplementary Figure 5**). We then injected retrograde AAV in the bPN to express Cre in all afferent areas to bPN. Subsequent injections with AAVs to express TVA and optimized G protein were made into the MDJ, followed by pseudotyped rabies virus one week later. In these experiments (**Figure 4A,** n=3 mice) we observed starter neurons in the MDJ that were GAD-positive (**Figure 4B**). As expected, in the bPN we could observe many axon terminals from starter neurons that contained labeling from the rabies virus, and were GAD-positive (**Figure 4C**). This confirmed that MDJ GAD-positive neurons indeed make contacts in the bPN. In the neocortex of these mice we could observe labeling of deep neocortical pyramidal neurons (**Figure 4D**). These results show that neurons in the MDJ that provide afferents to the bPN receive inputs from neocortex.

**Figure 4:**
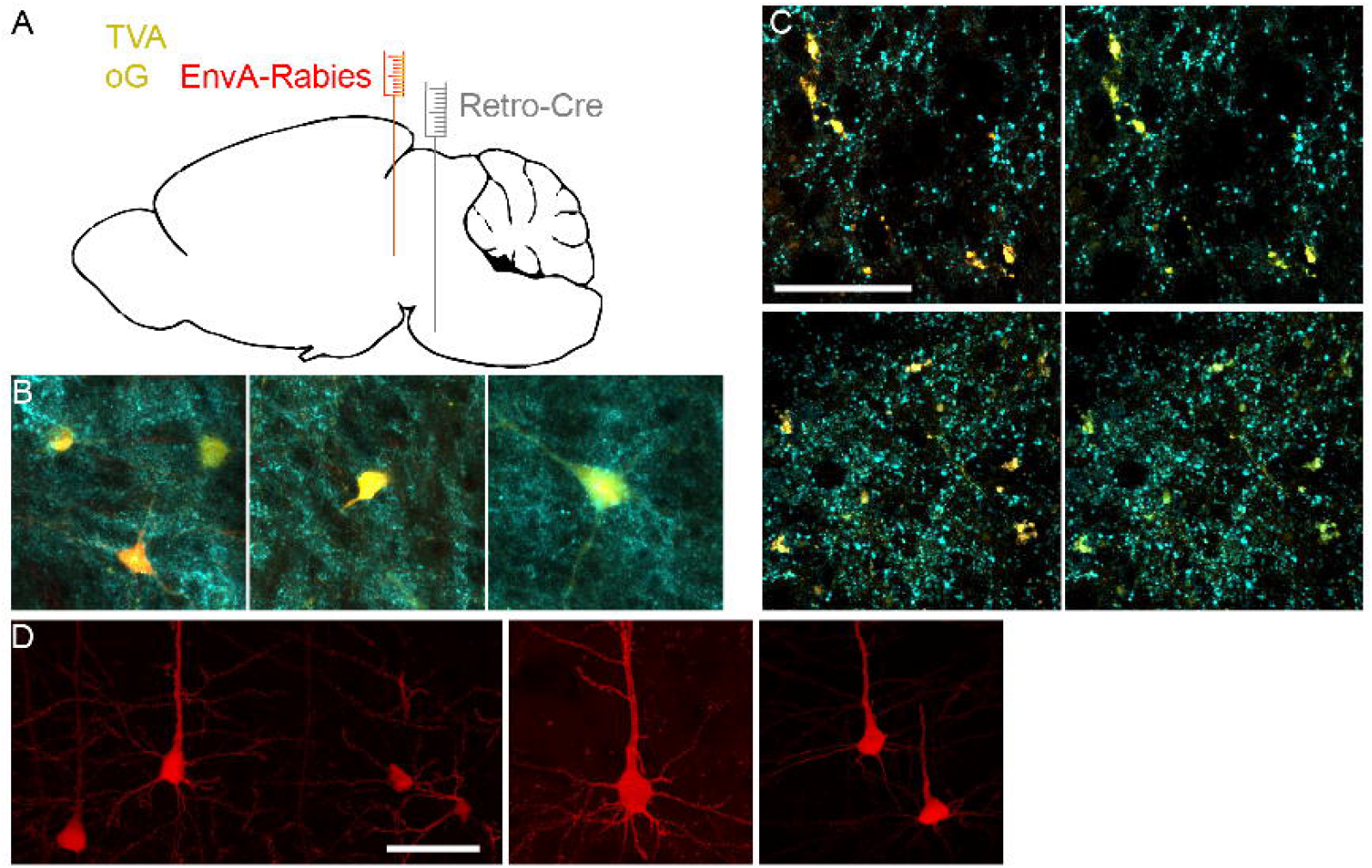
Rabies tracing of inputs to mesodiencephalic junction neurons that project to bPN. (A) Schematic representation of virus injections. (B) GABAergic rabies tracing starter neurons in mesodiencephalic junction were labeled with GFP from helper plasmids for rabies tracing. GAD67 staining (Cyan), Helper plasmids (Yellow), Rabies virus (Red). (C) GAD+ terminals from starter neurons can be observed in bPN. (D) Rabies tracing reveiled prominent deep layer labeling in neocortex. Scale bars are 50 μm.

## Discussion

We show that the bPN receives prominent inputs from neocortex and from MDJ. Using whole cell recordings and optogenetic stimulation we show that inputs from neocortex are glutamatergic and strongly depressing, while inputs from MDJ are GABAergic, but show remarkably little short-term depression. Interestingly, M1 and MDJ inputs do not interact at the single neuron level, nor via a network of synaptic connections in the bPN, and thus represent separate streams of information through the bPN, Finally, using Rabies-virus tracing we show that MDJ neurons that project to the bPN receive prominent input from neocortical output neurons. Thus, our results show and characterize an unknown connection from neocortex to bPN, which could provide sign-inversed inputs from neocortex to cerebellar granule cells.

### The source of inhibitory afferents to the bPN

The source of inhibition to the bPN has been unclear for a long time. Several afferent nuclei such as the zona incerta, anterior pretectal nucleus, cerebellar nuclei, prerubral area, the medullary formation (Border et al. 1986) and even neurons in the bPN itself (Border and Mihailoff 1985, 1990; Brodal et al. 1988) have been suggested to provide GABAergic inputs. In functional studies, inhibition of bPN neurons has been observed during behavior (Guo et al. 2021), and inhibition in bPN neurons has been observed in vitro after stimulation of the cerebral peduncle and the tegmentum (Möck et al. 1997). Indeed, in our present experiments, we observed prominent GABAergic inputs from MDJ, a group of neurons located in the tegmentum that receives prominent inputs from neocortex (De Zeeuw et al. 1998; Kubo et al. 2018). Furthermore, we have found no evidence that local interneurons are present in the bPN, nor that any neuron in the bPN makes local connections (see also Möck *et al*., 2006). GABAergic inhibition to bPN neurons therefore seems to be completely extrinsic. However, we can’t exclude that other sources than MDJ inputs may contribute to inhibition in the bPN

### bPN network architecture

The bPN are considered to integrate incoming motor and sensory information from the neocortex at the single cell level (Potter et al. 1978). Indeed, some neurons in the bPN respond only to movement whereas others are responsive to multiple modalities such as movement and cue (Guo et al. 2021), but this might also represent integration in neocortex. Although the precise extent of convergent streams in the bPN remains an important unanswered question, based on anatomical tracing data it is suggested that convergence of excitatory afferents from different regions to single bPN neurons is likely (Mihailoff et al. 1988; Lee and Mihailoff 1990; Schwarz and Thier 1999; Leergaard 2003; Leergaard and Bjaalie 2007) It is all the more striking then that we did not find any convergence of excitatory cortical and inhibitory MDJ inputs in the bPN. Although we cannot unequivocally rule out the presence synaptically connected bPN neurons, we expect that such connections would be too sparse to hold a substantial functional role.

### Difference in short-term plasticity between afferents

We show that MDJ GABAergic and glutamatergic cortical inputs are markedly different in their short-term plasticity. Cortical inputs show clear synaptic depression across all tested frequencies, which is particularly strong at relatively high frequencies. Conversely, MDJ inputs undergo little synaptic plasticity at all except for slight depression towards the end of a pulse train at higher stimulation frequencies. These differences are important, since synaptic depression or potentiation plays an important role in shaping the activity of neurons (Silver 2010). Layer 5 neurons provide the output from neocortex to bPN (Tervo et al. 2016), and respond with changes in firing rate up to 50Hz during movement (Park et al. 2021; Guo et al. 2021). According to our electrophysiological data, signals from neocortex below 50Hz will undergo limited short-term depression, and thus can be faithfully transmitted to bPN neurons. Indeed, during reaching, bPN neurons show modulations of their firing rates in line with activity in layer 5 of neocortex (Guo et al. 2021). Interestingly, GABAergic inputs from the MDJ undergo very little short-term depression. It is important to note that our estimates of short-term depression were obtained from optogenetic stimulations, which might induce artificial synaptic depression (Jackman et al. 2014). Indeed, Chrimson, a slower variant of channelrhodopsin showed more pronounced depression than Chronos, a fast variant (Klapoetke et al. 2014). Still, for frequencies up to 20Hz, both Chrimson and Chronos-mediated stimulation yielded very comparable results, indicating that depression in glutamatergic connections from neocortex probably are not an artifact from optogenetic stimulation.

### Potential roles of inhibitions in the bPN (gating vs. gain setting vs. enrichment)

It remains unclear by which mechanism inhibition in the bPN contributes to voluntary motor control. Although our findings do not decisively point to one single mechanism, we are able to rule out several hypotheses. Our rabies tracings suggest that GABAergic MDJ afferents to the bPN could be recruited by cortical activation. This possibly explains why optogenetic stimulation of the cortex either induces an increase or a decrease in the firing rate of bPN neurons (Guo et al. 2021). Although complete optogenetic silencing of the bPN disturbs movement (Wagner et al. 2019; Guo et al. 2021), our data show that inhibition from the MDJ specifically avoids bPN neurons that receive inputs from motor cortex. Therefore, we consider it unlikely that feedforward inhibition from the MDJ serves as a general gating mechanism (Crowley et al. 2009; Geborek et al. 2013). Furthermore, the phasic nature of GABA signaling in the bPN suggests a timing-dependent mechanism rather than continuous response gain adjustment (Silver 2010). It is therefore likely that the MDJ provides the MDJ with a negative signal based on neocortex inpus to MDJ. In this arrangement the bPN would transmit one direct, positive signal based on direct corticopontine inputs, and one negative signal based on cortico-MDJ-pontine inputs. This would greatly enrich the inputs that are provided to the input layer of the cerebellar cortex, which would support cerebellar learning (Chabrol et al. 2015; Cayco-Gajic et al. 2017; Straub et al. 2020).

## Acknowledgements

This work was supported by an NWO vernieuwingsimpuls VENI grant 016.Veni.171.056. The authors thank J.C. Lodder, A.J. Timmerman, T.S. Heistek and J Wortel for their excellent technical assistance. The authors also thank S. Abirashid, V Zucconi Galli Fonseca and J.K.W. de Vries for help with experiments and F.J. Meye and R.A.H. Adan for help with designing the rabies virus experiment. EnvA-complemented rabies virus (SAD-dG-tdTomato) was a generous gift from K.K. Conzelmann (DFG project-ID 118803580-SFB 870). pAAV-Syn-Chronos-GFP and pAAV-syn-ChrimsonR-tdT were a generous gift from Edward Boyden.

## Author Contributions

L.W. designed the study. A.K. performed and analyzed electrophysiological experiments. L.W. performed and analyzed anatomical experiments. All authors checked data analysis. A.K. and L.W. wrote the manuscript. All authors critically revised the manuscript.

## Supplementary figures

**Supplementary Figure 1: Injection locations for retrobead tracing from bPN**.

Two mice were injected with retrobeads in their bPN (top and bottom). Shown are the locations where tracer was injected with the atlas superimposed.

**Supplementary Figure 2: Optogenetic stimulation of ChrimsonR and Chronos expressing MDJ and M1 afferents**

MDJ inputs (left) were stimulated via Chronos (blue) or Chrimson (Orange) in separate experiments. When stimulating with Chrimson more pronounced synaptic depression can be observed compared with Chronos. Especially at higher frequencies (>50Hz) and at the end of the train (second column) this is more pronounced. M1 inputs (right) were stimulated in a similar manner via Chronos and Chrimson, resulting in comparable results.

**Supplementary Figure 3: GAD staining and GAD-GFP mice show the absence of GABAergic neurons, but the presence of GABAergic fibers In bPN**

(A) GAD67 staining of sections of mice show that there are no GABAergic neurons in the bPN of mice. Compare bPN with areas with known prominent GABAergic neurons (IPR, IPC, MM). RtTg: Reticulotegmental nucleus of the pons; IPC: Caudal subnucleus of the Interpeduncular Nucleus; IPR: Rostral subnucleus of the Interpeduncular Nucleus; MM: Medial Mammillary Nucleus. (B) Enlarged view of bPN showing prominent GABAergic fibers. (C) GAD-GFP mice show that there is absence of GABAergic neurons in bPN, but GABAergic fibers can be distinguished.

**Supplementary Figure 4: Electrophysiological characterization of passive membrane properties of MDJ and M1-receiving bPN neurons**

Membrane resistance, membrane time-constant and capacitance show no systematic difference between groups of MDJ and M1 input-receiving neurons. Horizontal lines represent single neurons, box plot shows median, 25^th^ and 75^th^ percentile. Bars represent 10^th^ and 90^th^ percentile.

**Supplementary Figure 5: Rabies tracing of inputs to bPN**.

(A) Schematic representation of injection sites for AAV-Cre (grey), AAVs with TVA and glycoprotein (yellow) and EnveA-Rabies (red). (B) After injection of rabies virus in bPN, prominent labeling of deep layer pyramidal neurons was observed, scale bar 50 μm. (C) In bPN starter neurons (Yellow) could be found together with afferents to bPN (Red). GAD67 staining (Cyan) indicates that some afferents, but none of the starter neurons were GABAergic. (D) Enlargement of the area shown with a white box in (C) indicating overlap between GAD staining and some afferents, but none of the starter neurons. Shown is a summed stack (left column) through with orthogonal view (right column) at the location indicated with the black arrowheads in the bottom stack. Scale bar 5 μm. (E) In cerebellar cortex, clear mossy fibers from bPN starter neurons could be found. Scale bar 50 μm. (F) In MDJ neuron could be observed that were positive for rabies virus, and positive for GAD67, indicating a GABAergic input from MDJ to bPN.

