## Supplementary figures and images for "Mesodiencephalic junction Gabaergic inputs are processed separately from motor cortical inputs in the basilar pons"

### Supplemental figures

Injection 1

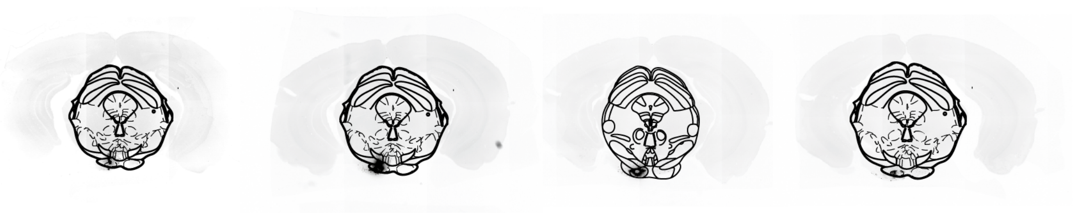

Injection 2

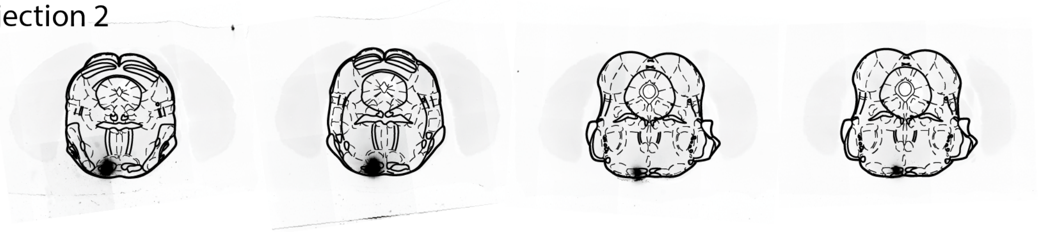

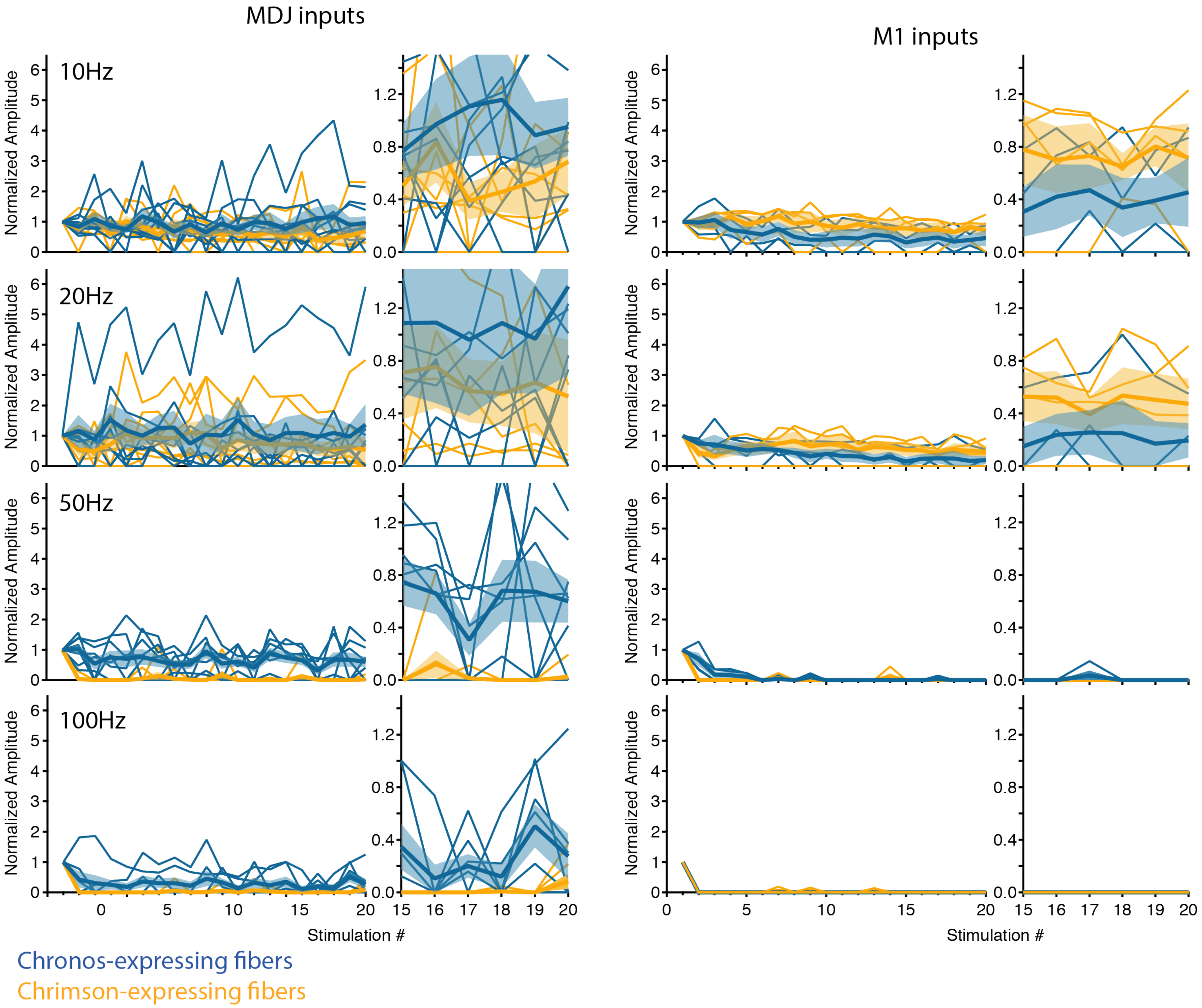

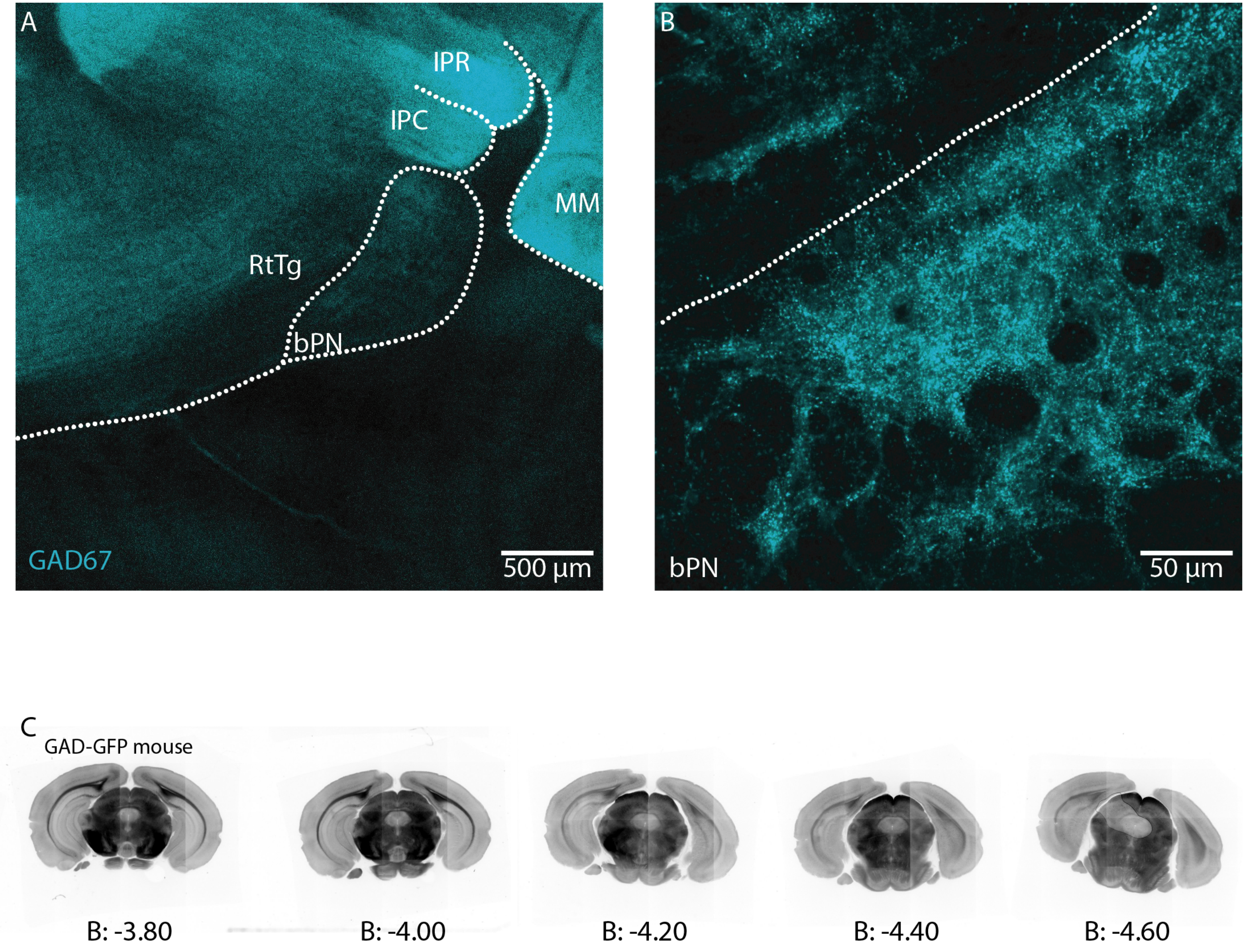

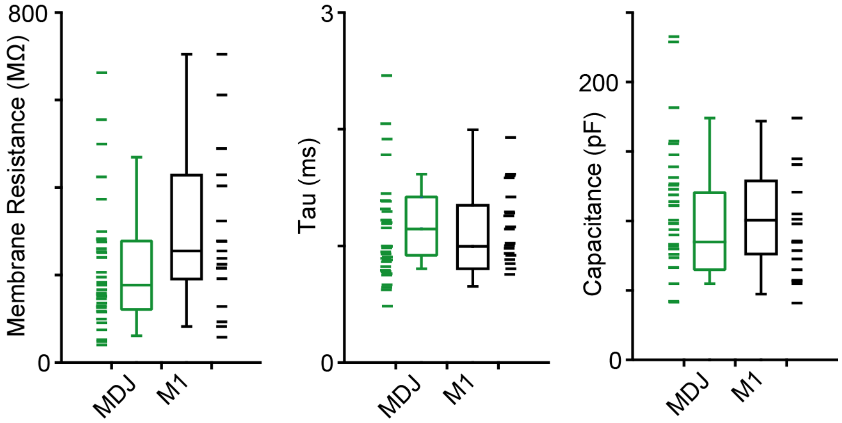

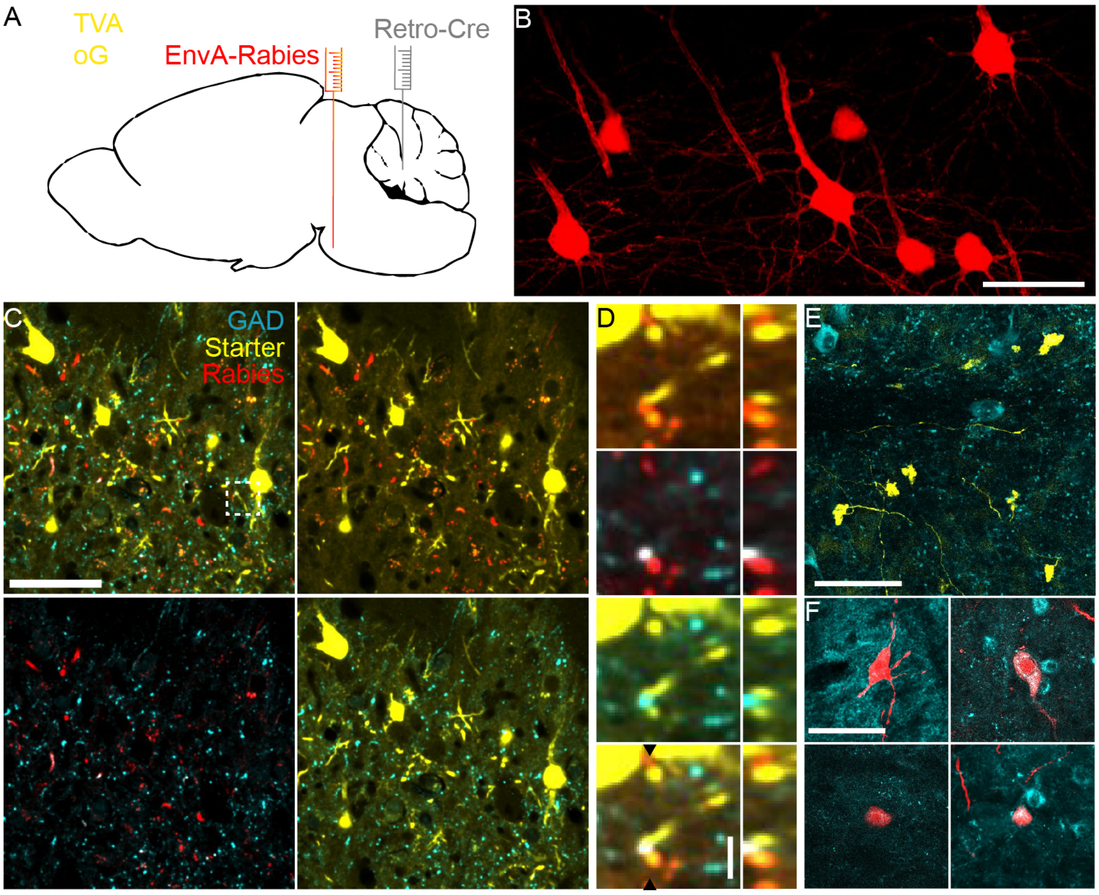
